# Neuroprotective Effects of Bioactive Molecules Derived from Tobacco as Potential Therapeutic Candidates for Alzheimer Disease

**DOI:** 10.1101/2024.03.20.585935

**Authors:** Ceheng Liao, Meng Li, Zhu Li, Meixia Wang, Qiyuan Peng, Zhouhai Zhu, Hong-Hui Wang, Ying Guan

**Author notes:** Corresponding author Hong-Hui Wang, Ying Guan.

## Abstract

Neurodegenerative diseases are significant global health challenges, particularly with an aging population. While tobacco is traditionally linked to health risks, recent studies suggest it may contain compounds beneficial for neurodegenerative conditions. Herein, we explore the potential of bioactive compounds in tobacco as neuroprotective agents for Alzheimer’s disease (AD). Using genetic engineering, we developed a novel approach with neural progenitor cells (NPCs) derived from embryonic stem cells, equipped with an NF-κB reporter system to screen tobacco extracts. Our screenings identified three compounds with significant inhibitory effects on NF-κB activation, a key mediator of neuroinflammation in AD. Among these, rutin exhibited profound neuroprotective effects in an NPC damage model induced by Amyloid-β25-35, reducing apoptotic cell death, enhancing cellular proliferation, and activating critical survival signaling pathways. This modulation underlies rutin’s anti-inflammatory and neuroprotective activities. Together, our findings support the potential of tobacco-derived compounds in AD therapy and lay the foundation for further exploration of their pharmaceutical value.

## Introduction

Neurodegenerative diseases, marked by their progressive and debilitating nature, are becoming a formidable challenge to global health (1). Currently, they afflict approximately 50 million people worldwide, with Alzheimer and Parkinson being the most prevalent neurodegenerative symptoms(2, 3). The aging as a risk factor increase global demographic in patients(4). Particularly, Alzheimer’s disease, is notorious for its profound memory loss, cognitive impairment, and behavioral changes [3]. The complexity of these neurodegenerative diseases, which involves diverse theories ranging from cholinergic deficits to genetic factors, adds to the difficulty in developing effective treatments (2, 3). Unfortunately, existing therapies are limited and often fail to address the diseases comprehensively, underscoring an urgent need for groundbreaking therapeutic strategies (1, 5).

Over the past several decades, the cultivation and utilization of tobacco (Nicotiana tabacum) have predominantly been driven by its nicotine content, which is associated with a multitude of health risks(6, 7). However, as scientific inquiry advances, a more nuanced understanding of tobacco’s pharmacological potential has revealed a plethora of compounds beyond nicotine that may confer therapeutic and health benefits(8, 9). This is particularly relevant in the field of neuroscience, where there is a growing demand for therapeutic agents that can mitigate or treat AD and Parkinson’s diseases (7, 10). Epidemiological studies have hinted at a correlation between smoking and a lower incidence of AD, suggesting that certain active components in tobacco may exert neuroprotective effects(8, 11). Nicotine, a principal constituent of tobacco, has been shown to ameliorate neural damage and elicit anti-inflammatory responses in glial cells(12). Furthermore, non-nicotine compounds present in tobacco, including rutin, chlorogenic acid, and solanine, have been recognized for their antioxidative, anti-inflammatory, and neuroprotective properties(6, 13-15). These insights offer a fresh perspective on assessing the economic value and potential applications of tobacco. In this context, tobacco producer becomes significant potentials for drug screening specifically for the treatment of neurodegenerative diseases, given its abundant and largely untapped tobacco leaf resources (6, 14, 16). Nevertheless, there remains a research gap in fully understanding the complex cellular processes of the human brain in relation to these compounds.

The pivotal role of neural progenitor cells (NPCs) in the protection and regeneration processes is essential for treating neurodegenerative diseases like AD(17, 18). NPCs hold the potential to restore cognitive function and neuronal integrity by differentiating into various neural cell types, replacing lost neurons, and integrating into existing neural networks(17). AD is characterized by a progressive decline in cognitive functions, leading to significant impairment in daily living. The amyloid cascade hypothesis posits that the accumulation of amyloid-β (Aβ) species, including monomers, oligomers, fibrils, and plaques, is a central event in the AD pathogenesis (19). The amyloid cascade hypothesis emphasizes the primacy of Aβ species in AD pathogenesis, while the emerging understanding of neuroinflammation highlights the importance of the immune response in the disease’s progression (20). Inflammatory processes, initiated by activated microglia and perpetuated by a cascade of cytokines and chemokines, can lead to synaptic dysfunction, neuronal damage, and ultimately, neurodegeneration(21). Therefore, identifying bioactive molecules that regulate inflammation and protect NPCs is crucial for neurodegenerative disorder treatment. These molecules may inhibit pro-inflammatory cytokines and bolster the brain’s anti-inflammatory responses, enhancing neural health. By mitigating inflammation, they improve the environment for NPCs, potentially restoring cognitive function and combating diseases like AD.

In this study, a human neural progenitor cell (NPC) model was developed to explore tobacco’s therapeutic potential for neurodegenerative diseases like Alzheimer’s. Utilizing genetic engineering, NPCs was engineered with an NF-κB reporter system and were used to identify effective tobacco extracts against neurodegeneration. Amyloid-β25-35 was shown to cause cytotoxicity, with rutin notably reducing apoptosis and promoting anti-apoptotic mechanisms. The study also investigated rutin levels in different tobacco processing methods, finding lower concentrations in air-dried compared to flue-cured tobacco. This research underscores tobacco-derived molecules, especially rutin, as promising for further pharmaceutical investigation into neurodegenerative treatments.

## Methods

### Cell Culture and Differentiation

Human neural stem cells (NSCs) were obtained from Cellway Bio and cultured in NSC proliferation medium supplemented with epidermal growth factor (EGF) and basic fibroblast growth factor (bFGF) as per the manufacturer’s instructions. NSCs were maintained in a humidified incubator at 37°C with 5% CO2.

### NF-κB Reporter Cells for Evaluation of Neural Inflammation

The NF-κB reporter plasmid (Lenti-NFκB-mCherry), containing the mCherry gene under the control of a minimal promoter with five tandem NF-κB binding sites, was co-transfected with the viral packaging vectors pCMV-VSV-G and pCAG-dR8.9 into HEK 293T cells using Lipofectamine 3000 according to the manufacturer’s protocol. Lentiviral particles were produced and used to infect human neural stem cells. Positive clones were selected using limited dilution and expanded for further experiments, resulting in a stable NF-κB reporter cell line, NPC-NF-κB-mCherry. These cells were seeded in 96-well plates and pretreated with various concentrations of tobacco-derived bioactive molecules for 2 hours before Aβ25-35 treatment. After 24 hours, cells were imaged, and fluorescence intensity was quantified using a microplate reader.

### Aβ25-35 Preparation and Treatment

Aβ25-35 peptide was dissolved in sterile distilled water at a concentration of 1 mM and incubated at 37°C for seven days to allow aggregation. NPCs were treated with various concentrations of aggregated Aβ25-35 (0, 20, 50, and 100 µM) for 24 hours to determine cytotoxic effects using the CCK8 assay as per the provider’s guidelines.

### Extraction of Tobacco-derived Rutin using High-Performance Liquid Chromatography (HPLC)

Tobacco samples encompassing various curing processes—Initial Curing, Re-curing, and Air-curing—were procured from local suppliers, representing different tobacco species such as Red Da Tobacco and K326 Tobacco. The initial curing refers to the first stage of tobacco processing where the leaves are dried to initiate the chemical changes necessary for flavor development. Re-curing involves further drying and aging to enhance and refine the tobacco’s quality, while air-curing is a natural drying method that allows the leaves to dry over time in a ventilated space, preserving their natural characteristics. For the extraction of rutin, a well-established method was employed. The extracted samples were passed through a 0.22 µm syringe filter to remove particulates before undergoing High-Performance Liquid Chromatography (HPLC) analysis. The HPLC system was fitted with a C18 column, which is commonly used for the separation of compounds based on their polarity. The mobile phase used in the HPLC was a combination of solvent A, which is 0.1% formic acid in water, and solvent B, which is 0.1% formic acid in acetonitrile. This mixture was crucial for the efficient separation of rutin from other compounds present in the tobacco samples. The gradient elution profile was meticulously controlled, starting with 15% B for the first 10 minutes, increasing to 50% B by 30 minutes, decreasing back to 15% B by 35 minutes, and maintaining this percentage until the 40-minute mark. This dynamic change in the mobile phase composition helps to resolve complex mixtures and elute rutin effectively. The flow rate throughout the analysis was maintained at 1 mL/min, ensuring a consistent flow of the mobile phase through the column. The injection volume for each sample was 20 µL, which is a standard volume for HPLC injections to ensure accurate and reproducible results. Rutin detection was achieved using a UV detector set at a wavelength of 360 nm, which is specific for the detection of flavonoids like rutin. The content of rutin in the samples was quantified by comparing the peak areas to a calibration curve generated using known concentrations of rutin standards. This method allows for the accurate determination of rutin content in various tobacco samples, which is essential for understanding the potential health effects and quality of tobacco products.

### Western Blot Analysis

Rutin was dissolved in dimethyl sulfoxide (DMSO) and further diluted in culture medium to achieve the desired concentrations. NPCs were pretreated with rutin for 2 hours before Aβ25-35 treatment. Cells were lysed in RIPA buffer supplemented with protease and phosphatase inhibitors. Protein concentration was determined using the BCA Protein Assay Kit. Equal amounts of protein were separated by SDS-PAGE and transferred to PVDF membranes. Membranes were probed with primary antibodies against NF-κB (1:500), pro-caspase-3 (1:1000), cleaved-caspase 3 (1:1000), Bcl-2 (1:1000), and tubulin (1:5000), followed by HRP-conjugated secondary antibodies (1:5000). Protein bands were visualized using an enhanced chemiluminescence (ECL) detection system, and densitometry analysis was performed using ImageJ software.

### Immunofluorescence Assay

For assessing neuronal differentiation, NPCs were fixed with 4% PFA, permeabilized with 0.1% Triton X-100, and blocked with 1% BSA. They were incubated with a Tuj-1 primary antibody (1:500) overnight at 4°C, then with a goat anti-rabbit secondary antibody conjugated with Alexa Fluor 567. Nuclei were stained with DAPI for blue fluorescence. To evaluate rutin’s neuroprotection, NPC-NF-κB-mCherry cells were pre-treated with 1 µg/mL rutin for 2 hours before exposure to 50 µM Aβ25-35 peptide. After fixation with 4% PFA and permeabilization with 0.1% Triton X-100, cells were blocked with 1% BSA and incubated with a rabbit anti-human Ki67 antibody (1:100) for 1 hour, followed by a goat anti-rabbit secondary antibody conjugated with Alexa Fluor 567. Nuclei were again stained with DAPI.

### Statistical Analysis

All experiments were performed in triplicate, and data are presented as mean ± standard deviation (SD). Statistical significance was determined by unpaired two-tailed Student’s t-test, and *p□< □ 0.05 and **p<0.001 was considered to be statistically significant. ‘NS’ indicates not significant.

## Results

To construct a human-derived NSC line capable of visualizing the inflammatory responses, we developed a novel lentiviral vector named Lenti-NF-κB-mCherry for the establishment of a stable NSC line. In the gene construction experiment, we successfully cloned the regulatory binding sequences of the NF-κB protein as part of the promoter into the mCherry reporter gene vector, which was confirmed by sequencing (**Figure S1**). We then used the lipofection method to co-transfect this vector with the viral packaging vectors pCMV-VSV-G and pCAG-dR8.9 into HEK 293T cells for lentiviral packaging. The successful expression of the activated mCherry protein, indicated by red fluorescence, was observed in these cells (**Figure S2**). The genetic engineering process was based on our previously established NSC line, which is optimized for long-term culture with the ability to undergo over 50 passages in the laboratory and differentiate into neurons and glial cells(22). we infected human-derived NSCs with a lentivirus derived from Lenti-NF-κB-mCherry-transduced HEK 293T cells. Through the application of limited dilution, we successfully isolated and expanded positive clones, leading to the establishment of a stable NF-κB reporter cell line named NPC-NF-κB-mCherry. This cell line characteristically shows minimal basal fluorescence, indicating a low level of NF-κB activity, in the absence of inflammatory stimuli (**Figure S3**). This stable cell line retains the ability of NSCs to form spheres and differentiate into Tuj-1 positive neurons (**Figure S4**).

Using the established NPC-NF-κB-mCherry cell line, we investigated the assessment of neuroinflammatory responses. Compared to control healthy NSCs, the addition of the pro-inflammatory TNFα (1nM) significantly increased red fluorescence in the NPC-NF-κB-mCherry cell line (**Figure 1a**). The intensity of red fluorescence increased over time, reaching 27.2 times that of the untreated group after 24 hours (**Figure 1b**). Furthermore, we used immunoblotting to detect the expression levels of NF-κB protein. The results showed that NF-κB protein expression also increased with extended treatment time (**Figure 1c**). These findings indicate that TNF stimulation of NF-κB expression levels can be characterized by live-cell fluorescence imaging, allowing for the quantitative assessment of neuroinflammatory signals. The fluorescent reporter system we constructed enables the quantitative evaluation of cellular inflammatory levels.

**Figure 1:**
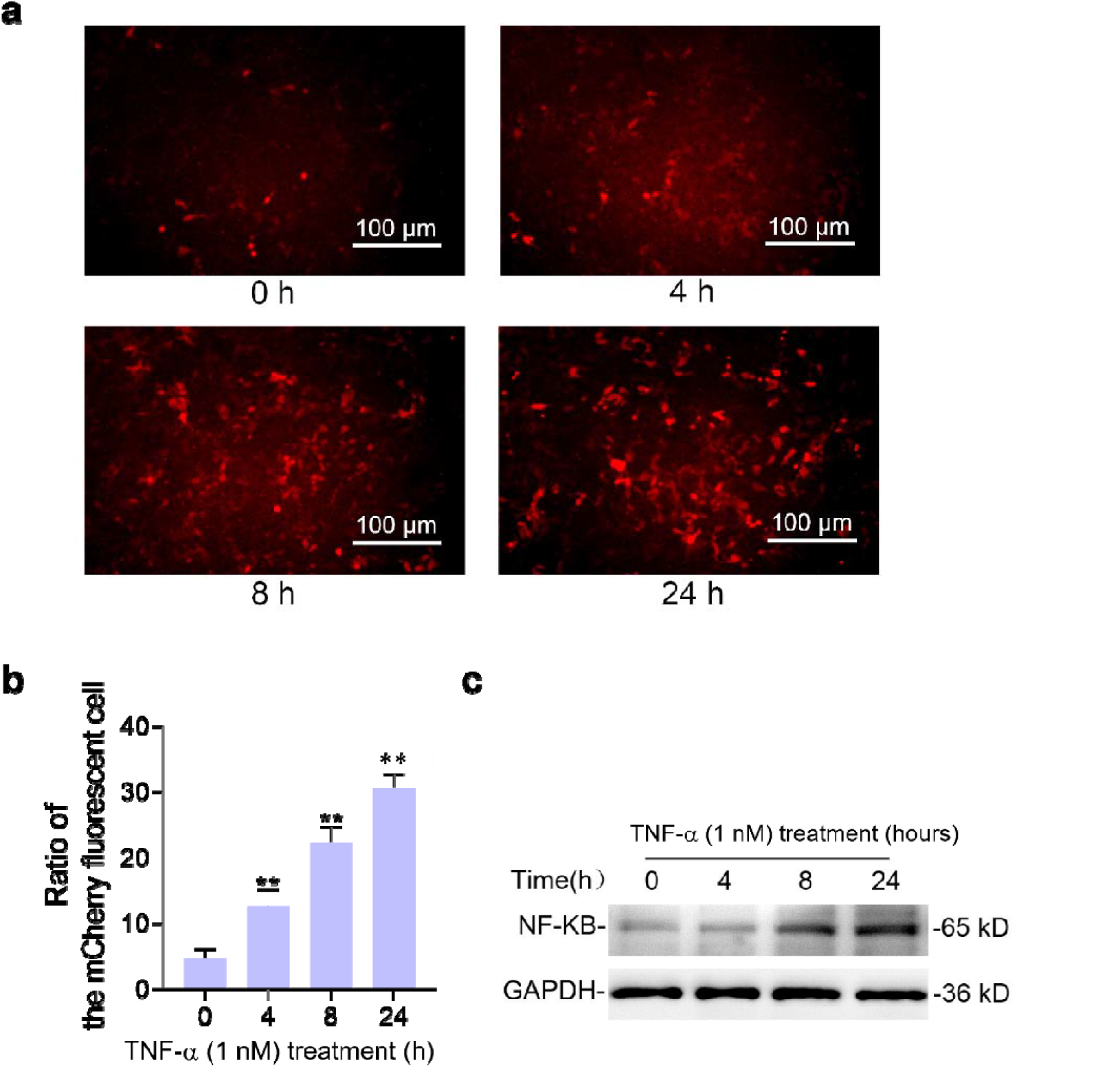
Construction of NPC-NF-κB-mCherry cells for visualizing neural inflammation. a. Live-cell imaging captures of NPC-NF-κB-mCherry cells at key time intervals (0, 4, 8, and 24 hours) post-TNF-α treatment. The images indicate the translocation of NF-κB to the nucleus for transcriptional regulation, indicated by an increase in nuclear mCherry fluorescence, which serves as a visual marker for neuroinflammation. The scale bar is 100 μm. b. Graphical representation of the quantitative fluorescence analysis from the live-cell imaging in (a). Data points represent the mean ± standard deviation (S.D.) of five independent experiments. Statistical significance is indicated by *p<0.05 and **p<0.001. c. Western blot analysis quantifies NF-κB protein levels in NPC-NF-κB-mCherry cells at different time points following TNF-α exposure. The accompanying graph illustrates the densitometry analysis of NF-κB protein levels relative to the loading control, providing a quantitative assessment of protein expression changes over time.

This cell line can be utilized to effectively assess the neuroinflammatory effects in an in vitro cell model of AD. We treated NPC-NF-κB-mCherry cells with β-amyloid peptide (Aβ25-35), whose aggregation and cytotoxicity are used to evaluate potential AD therapeutic drugs(19). We found that treatment with 50 µM Aβ25-35 for different durations resulted in a noticeable mCherry red fluorescence signal after 12 hours, with saturated fluorescence observed at 24 hours, indicating significant activation of NF-κB activity (**Figure 2a**). Subsequently, we validated the association of cell viability with the decrease in Bcl-2 levels using the CCK-8 assay. The addition of 50 µM Aβ25-35 significantly affected the viability of the NPC cells (**Figure 2b**), leading us to continue using this concentration to construct the cell damage model. We assessed the impact of different concentrations of Aβ25-35 on cell proliferation and found that cell viability decreased with increasing peptide concentration after 24 hours, consistent with the role of Aβ25-35 in the pathogenesis of AD (**Figure 2b**). Furthermore, the anti-apoptotic role of Bcl-2 is pivotal as it prevents programmed cell death by blocking the release of cytochrome c from mitochondria, a key event in the activation of caspases, which are the enzymes responsible for carrying out the cell’s apoptotic program(23). We next quantified the Bcl-2 expression using Western blot analysis and our findings revealed that exposure to Aβ25-35 led to a reduction in Bcl-2 levels in a manner that was dependent on the concentration of the peptide (**Figure 2c**). This decrease in Bcl-2 expression suggests a potential shift towards a pro-apoptotic state in the cells, which could be a contributing factor to the neurodegenerative processes observed in Alzheimer’s disease models(24).

**Figure 2:**
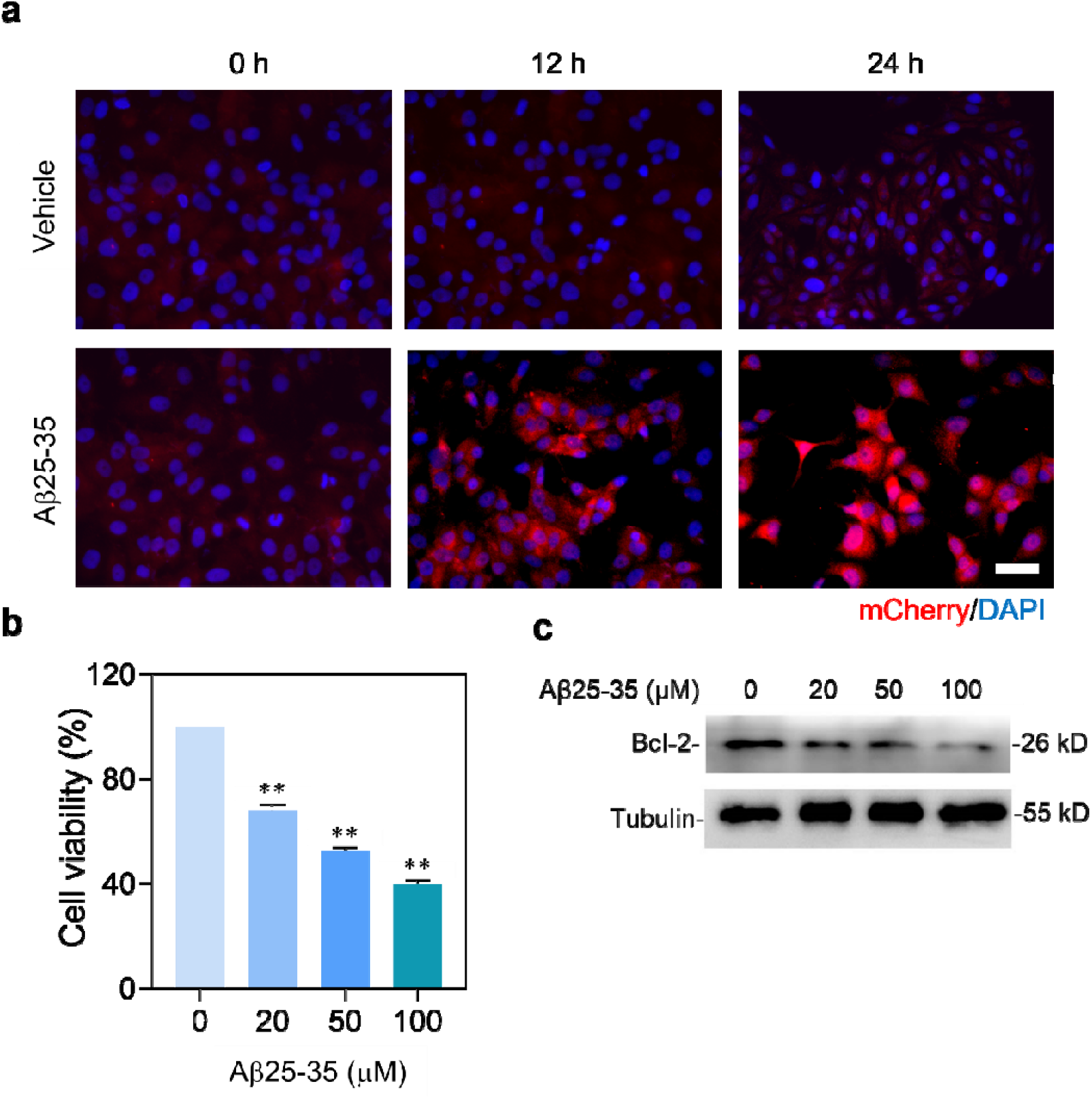
Establishing an inflammatory response and neuronal injury model in NPC-NF-κB-mCherry cells induced by Aβ25-35. a. Representative fluorescence images of NPC-NF-κB-mCherry cells treated with Aβ25-35 and vehicle at various time points (0, 12, and 24 hours), with nuclei visualized by DAPI staining. The scale bar corresponds to 20 μm, highlighting the temporal progression of neuroinflammation as indicated by the mCherry fluorescence. b. Cell viability assay results using the CCK8 method in NPC-NF-κB-mCherry cells exposed to different concentrations of Aβ25-35 for 24 hours. Data are expressed as the mean ± standard deviation (S.D.) (n = 5), with *p<0.05 and **p<0.001 denoting statistically significant differences, reflecting the dose-dependent effect of Aβ25-35 on cell viability. c. Western blot analysis determining Bcl-2 protein levels in NPC-NF-κB-mCherry cells treated with varying concentrations of Aβ25-35 to evaluate alterations in the expression of anti-apoptotic proteins, which may be indicative of the cells’ response to neurodegenerative stress.

Having established the NPC neuroinflammation and damage model based on Aβ25-35 aggregation, we further explored the protective effects of tobacco-derived bioactive molecules on neural progenitor cells. NPC-NF-κB-mCherry cells pre-treated with different tobacco extracts, including rutin, nornicotine, and chlorogenic acid(25). The results showed rutin and nornicotine could promote the healthier cell morphology relative to control groups and chlorogenic acid treatment (**Figure 3a**). These findings indicate rutin and nornicotine might exhibit potential neuroprotective effects, possibly due to the inhibition of pro-inflammatory signals, consistent with previous reports(26). Importantly, cell viability assays using the CCK-8 assay after 24 hours of pre-treatment with these molecules, followed by Aβ25-35 exposure, revealed that rutin and nornicotine significantly enhanced cell viability (Figure 3b), aligning with observations from optical microscopy. Furthermore, we examined the levels of the anti-apoptotic protein Bcl-2 and found that rutin and nornicotine rescued the decrease in Bcl-2 protein levels caused by Aβ25-35, indicating that these molecules may act through the Bcl-2-mediated anti-apoptotic pathway (**Figure 3c**). Relative to other molecules, rutin showed the strongest ability to enhance cell viability and rescue Bcl-2 protein levels, highlighting its significant role in neuroprotection.

**Figure 3:**
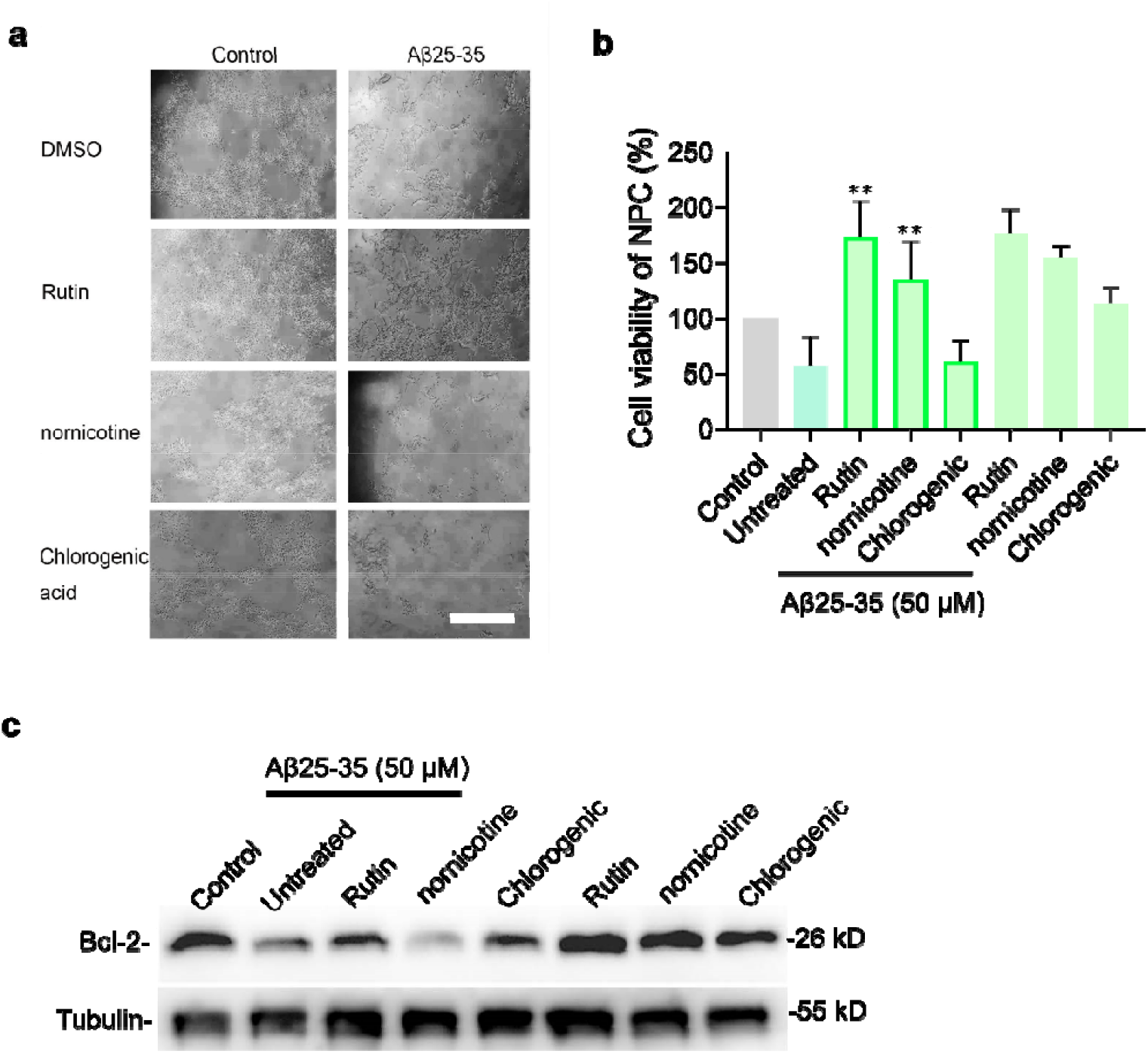
Neuroprotective effects of tobacco-derived bioactive molecules in an Aβ25-35-induced Alzheimer’s disease model. a. Light microscopy images of NPC-NF-κB-mCherry cells pretreated with different tobacco-derived bioactive molecules (rutin, nornicotine, and chlorogenic acid at 1 µg/mL) for 2 hours, followed by treatment with Aβ25-35 (50 µM) and vehicle. The scale bar represents 200 μm., illustrating the potential of these molecules to enhance cellular activity. b. Cell viability assay using the CCK8 method in NPC-NF-κB-mCherry cells pretreated with the specified tobacco-derived molecules for 2 hours, then exposed to Aβ25-35 for 24 hours. Data are presented as mean ± standard deviation (S.D.) (n = 5), with *p<0.05 and **p<0.001 indicating statistical significance. c. Western blot analysis of Bcl-2 protein levels in NPC-NF-κB-mCherry cells pretreated with the specified tobacco-derived molecules for 2 hours, then exposed to Aβ25-35 (50 µM) for 5 hours. Tubulin serves as the loading control. These results shed light on the potential neuroprotective mechanisms of these bioactive molecules, possibly through the modulation of the anti-apoptotic protein Bcl-2.

Considering the potential of rutin as a bioactive molecule with, we systematically compared rutin content in tobacco samples prepared by different methods using high-performance liquid chromatography (HPLC) (**Figure 4a and S2**). We found that air-cured tobacco, which has a darker brown color compared to flue-cured tobacco, contains significantly lower levels of rutin, indicating a direct relationship between the oxidative colorants produced by rutin and tobacco browning. The K326 series of tobacco, after flue-curing, had the highest rutin content at 10.67 mg/g, while the Hong Da tobacco series had a higher rutin content of 8 mg/g after flue-curing, compared to 3.33 mg/g in the initial curing stage (**Table S1**). This analysis provides insights into the content of rutin in different tobacco leaves and preparation processes, offering a basis for optimizing tobacco preparation techniques and identifying sources of molecules with anti-neural damage activity.

**Figure 4:**
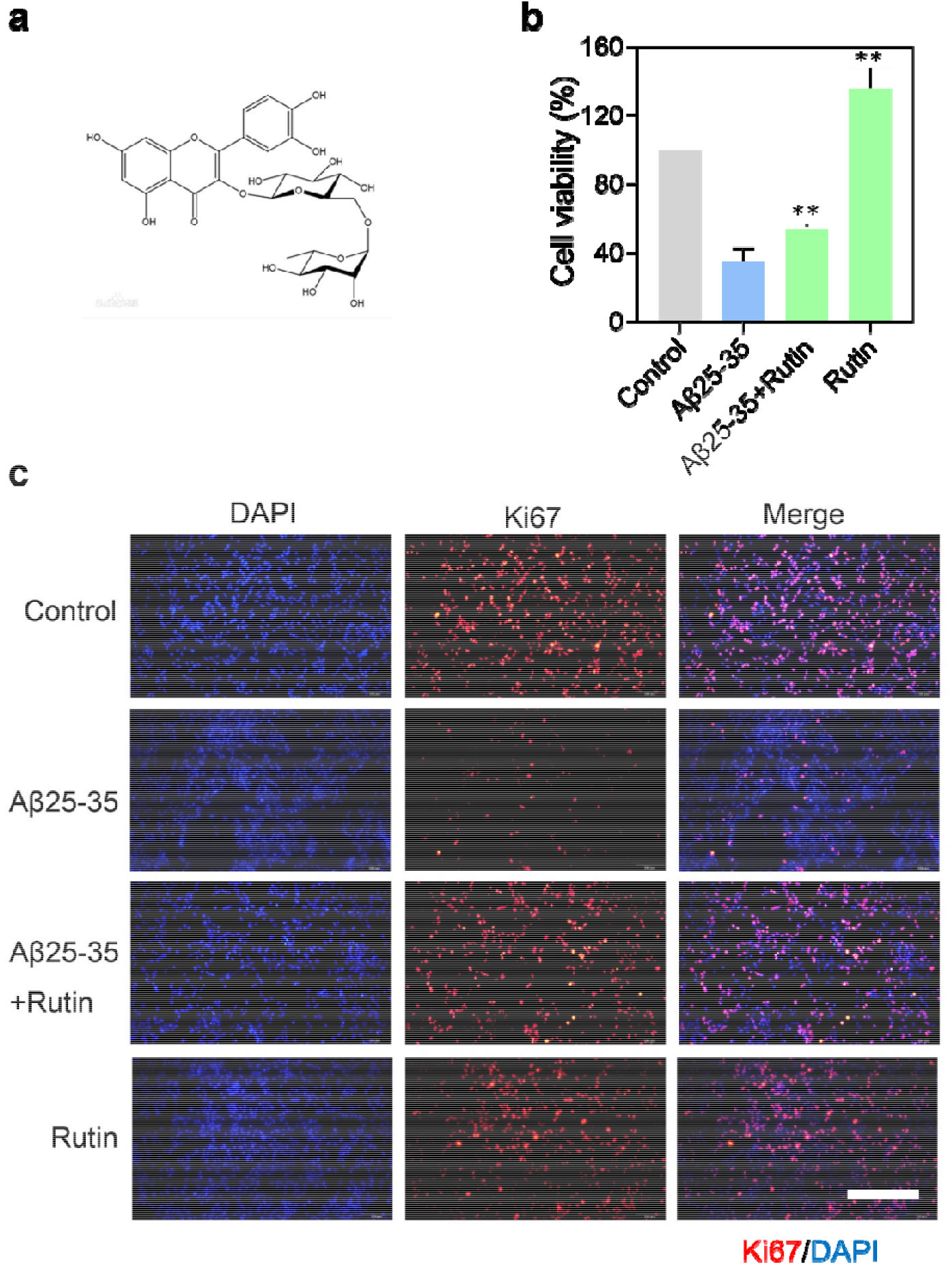
Rutin promotes proliferation of NPCs following Aβ25-35 injury. a. The chemical structure of rutin is depicted, highlighting its potential role in cellular protection and proliferation. b. Cell viability, as determined by the CCK8 assay, is assessed in NPC-NF-κB-mCherry cells pre-treated with rutin (1 µg/mL) for 2 hours and then challenged with 50 µM Aβ25-35 for 48 hours. Results are expressed as the mean ± standard deviation (S.D.) of five replicates, with * indicating p<0.05 and ** p<0.01 to denote statistical significance. c. Immunofluorescence staining for the proliferation marker Ki67 is performed in NPC-NF-κB-mCherry cells that were pre-treated with rutin (1 µg/mL) for 2 hours before exposure to 50 µM Aβ25-35. Nuclei are counterstained with DAPI. The scale bar is 200 μm.

We then investigated the pharmacological activity of rutin by examining its role in cell proliferation within the β-amyloid peptide-induced Alzheimer’s cell damage model. Pretreatment of cells with rutin significantly increased cell viability after 48 hours of Aβ25-35 exposure, to 1.36 times that of the disease model group (**Figure 4b**). Interestingly, pretreatment with rutin not only protected NPCs from damage but also enhanced their proliferation, indicating a role for rutin in the cell cycle. Ki67, a marker indicative of cell proliferation, is a nuclear protein associated with cell cycling, present during the active phases of the cell cycle (G1, S, G2, and M) and absent in quiescent (G0) cells(27). Immunofluorescence assays confirmed rutin’s role in promoting NPC proliferation, showing an increase in Ki67-positive cells following rutin treatment (**Figure 4d**).

To elucidate the molecular mechanism underlying rutin’s protective effect against Aβ25-35-induced apoptosis, we examined the expression levels of pro-caspase-3 and Bcl-2 by Western blot analysis. Caspase 3 cleavage is a critical step in the execution phase of apoptosis, the programmed cell death process. Activated caspase 3, a cysteine-aspartic protease, acts as the primary effector caspase that cleaves a variety of cellular substrates, leading to the dismantling of the cell and ultimately, cell death(28). We previously found that Aβ25-35 treatment significantly reduced Bcl-2 levels. We found rutin significantly rescued Bcl-2 in NPC cells, enhancing the protein expression of the anti-apoptotic factor Bcl-2, suggesting its potential to further inhibit cellular apoptosis (**Figure 5a and 5b**). In fact, rutin’s treatment effectively rescued the levels of cleaved caspase-3 induced by Aβ25-35 (**Figure 5a and 5c**). In neurodegenerative diseases, caspase 3 plays a significant role by promoting the apoptotic death of neurons, which contributes to the progressive neuronal loss observed in these conditions(29). Therefore, our findings suggest that rutin modulates anti-apoptotic Bcl-2 proteins to exert its neuroprotective effect, avoiding caspase 3 cleavage and apoptosis in Aβ25-35-treated NPCs. Additionally, rutin effectively promotes cell proliferation, enabling NPCs to resist damage caused by Aβ25-35 (**Figure 4b**). Overall, our findings demonstrate that rutin can effectively alleviate Aβ25-35-induced apoptosis in NPCs by inhibiting the activation of pro-caspase-3 and regulating the expression of Bcl-2.

**Figure 5:**
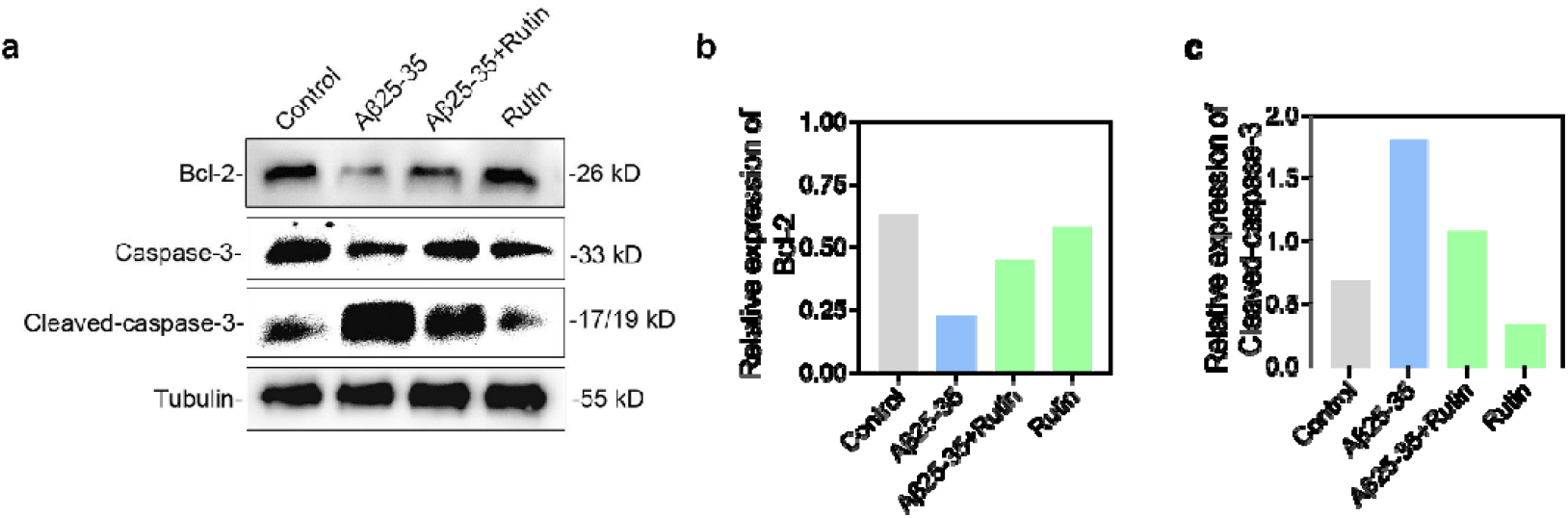
Rutin’s inhibitory effect on the apoptotic pathway in NPCs induced by Aβ25-35. a. Western blot analysis of the anti-apoptotic protein Bcl-2 and apoptotic key proteins, total and cleaved caspase 3, in NPC-NF-κB-mCherry cells. Cells were pretreated with rutin (1 µg/mL) for 2 hours followed by exposure to Aβ25-35 (50 µM) for 5 hours. Tubulin is used as a loading control. b. Quantitative analysis of Bcl-2 protein levels normalized to tubulin, as shown in Figure 5a. c. Quantitative analysis of the ratio of active (cleaved) caspase 3 to total caspase 3 in Figure 5a, demonstrating the potential of rutin to modulate the apoptotic pathway in response to Aβ25-35-induced stress.

## Discussion

Neurodegenerative diseases, such as Alzheimer’s and Parkinson’s, are increasingly prevalent as the global population ages, necessitating the development of novel therapeutic strategies(3, 30). We have harnessed genetic engineering to create the NPC-NFκB-mCherry cell line, an engineered human neural progenitor cell system that serves as a powerful tool for studying the neuroinflammatory processes associated with these diseases. This innovative cell line, equipped with a fluorescent sensor that reports NF-κB activity, allows for the real-time visualization of neuroinflammatory responses in vitro. By using this system, we have been able to model the complex neurodegenerative processes triggered by amyloid-beta peptides, akin to those seen in Alzheimer’s disease, and assess the potential neuroprotective effects of bioactive molecules derived from tobacco, such as rutin.

The creation and application of the NPC-NFκB-mCherry cell line mark a significant advancement in the study of neuroinflammatory mechanisms. This system enables researchers to observe and quantify NF-κB activity with high precision, providing valuable insights into the cellular responses to neurotoxic stimuli. By modeling the effects of amyloid-beta peptides, we can closely mimic the pathological conditions observed in Alzheimer’s disease, thereby offering a robust platform for testing therapeutic interventions. The use of bioactive molecules like rutin, derived from tobacco, has shown promising neuroprotective effects in our preliminary studies, highlighting the potential of natural compounds in mitigating neurodegeneration(15).

Looking ahead, the NPC-NFκB-mCherry cell line holds promise for integration into more complex tissue models, such as brain-like organoids (31). These three-dimensional cultures can provide a more physiologically relevant environment for validating the efficacy of potential therapeutics. The ability to visualize NF-κB activity within these organoids will offer high spatiotemporal resolution, enabling the tracking of neuroinflammatory processes across different tissue layers and cellular compartments. This integration will enhance our understanding of neurodegenerative disease mechanisms and facilitate the development of more effective treatments, ultimately contributing to improved outcomes for patients suffering from these debilitating conditions.

## Supporting Information

Methods and Materials, supplementary Tables and figures.

## Acknowledgement

We would like to acknowledge the Scientific Research Program from the Joint Institute of Tobacco and Health (Grant No. 2022539200340039).

## Author Contributions

All authors have given approval to the final version of the manuscript.

