## Supplemental Tables and Supplemental Figures for "Neuroprotective Effects of Bioactive Molecules Derived from Tobacco as Potential Therapeutic Candidates for Alzheimer Disease"

Corresponding author

**Table S1. Rutin content in different tobacco samples treated with various methods**

| Sample name | Peak Area (mAU) | Concentration (mg/ml) | Concentration (mg/g) |
|---|---|---|---|
| Initial Curing of Red Da leaves | 500 | 0.33 | 3.33 |
| Re-cured Red Da leaves | 1200 | 0.8 | 8.0 |
| Re-cured K326 leaves | 1600 | 1.07 | 10.67 |
| Air-cured K326 leaves | 50 | 0.03 | 0.33 |
| Air-cured K326 stalks | 5 | 0 | 0.03 |
| Rutin standard 1 | 60 | 0.04 | |
| Rutin standard 2 | 150 | 0.10 | |

Note: The last column "Concentration (mg/g)" is calculated based on the assumption that 1g of the sample was used for the concentration measurement. If the sample weight was different, this value needs to be adjusted accordingly.

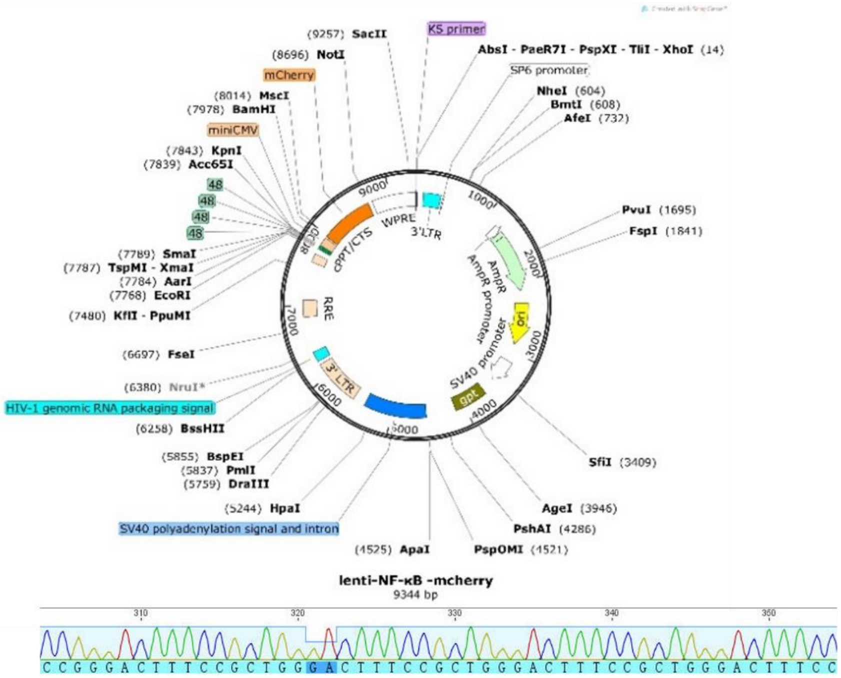


**Figure S1.** Schematic representation of the Lenti-NF-κB-mCherry plasmid (upper panel) and confirmation of successful NF-κB insertion into the mCherry promoter region via Sanger sequencing.

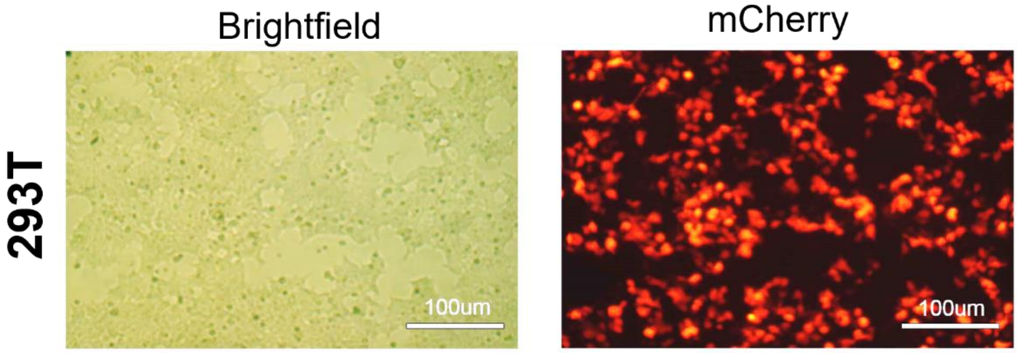


**Figure S2.** Fluorescent microscopic image of HEK 293T cells transfected with Lenti-NF-κB-mCherry plasmid and viral packaging vectors pCMV-VSV-G and pCAG-dR8.9, using Lipofectamine 3000 as per the manufacturer's protocol. The image was captured under a fluorescent microscope. Scale bar: 100 μm.

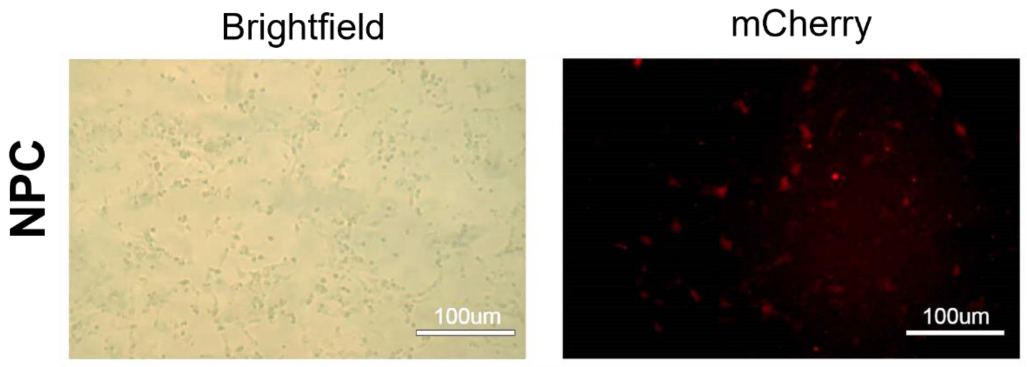


**Figure S3.** Light microscopic (left panel) and fluorescent microscopic image (right panel) of the human-derived NPC cell line infected with lentiviral particles from H293T cells transfected with Lenti-NF-κB-mCherry. The image was captured under a fluorescent microscope. Scale bar: 100 μm.

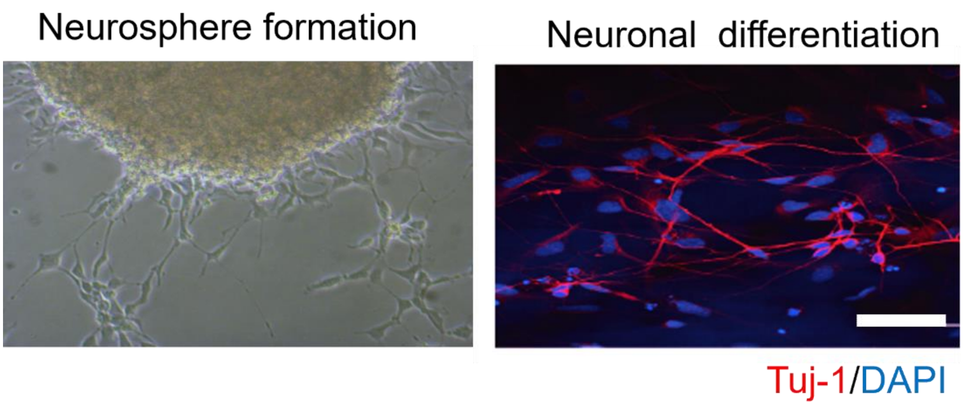


**Figure S4.** Light microscopy images of neurosphere formation (left panel) and immunofluorescence staining of NPCs following a neuronal differentiation protocol (right panel). Tuj-1 is labeled in red, and nuclei are counterstained with DAPI in blue. Scale bar: 50 μm.

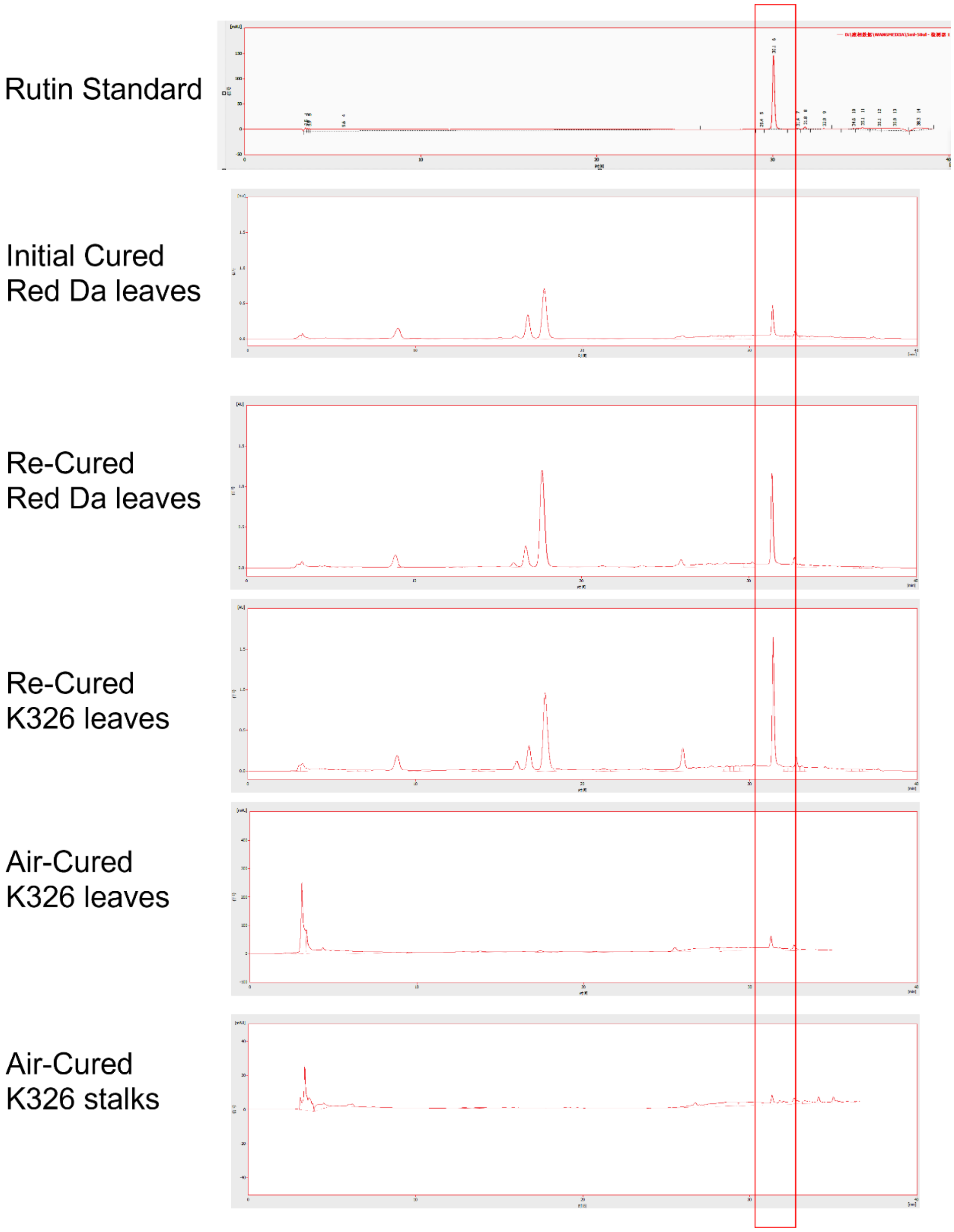


**Figure S5.** High-Performance Liquid Chromatography (HPLC) analysis of rutin content in different tobacco samples processed with various methods. The red box indicates the peak corresponding to rutin.
